# Transcriptional dynamics of colorectal cancer risk associated variation at 11q23.1 are correlated with tuft cell abundance and marker expression *in silico*

**DOI:** 10.1101/2022.03.29.485182

**Authors:** Bradley T. Harris, Vidya Rajasekaran, James P. Blackmur, Alan O’Callaghan, Kevin Donnelly, Maria Timofeeva, Peter G. Vaughan-Shaw, Farhat V. N. Din, Malcolm G. Dunlop, Susan M. Farrington

## Abstract

Colorectal cancer (CRC) is characterised by heritable risk that is not well understood. Heritable, genetic variation at 11q23.1 is associated with increased colorectal cancer (CRC) risk, demonstrating eQTL effects on 3 cis- and 23 trans-eQTL targets. We sought to determine the relationship between 11q23.1 cis- and trans-eQTL target expression and test for potential cell-specificity. scRNAseq from 32,361 healthy colonic epithelial cells was aggregated and subject to weighted gene co-expression network analysis (WGCNA). One module (blue) included 19 trans-eQTL targets and was correlated with *C11orf53* expression only. Following unsupervised clustering of single cells, the expression of 19 trans-eQTL targets was greatest and most variable in cluster number 11, which transcriptionally resembled tuft cells. 14 trans-eQTL targets were found to demarcate this cluster, 11 of which were corroborated in a second dataset. Intra-cluster WGCNA and module preservation analysis then identified twelve 11q23.1 trans-eQTL targets to comprise a network that was specific to cluster 11. Finally, linear modelling and differential abundance testing showed 11q23.1 trans-eQTL target expression was predictive of cluster 11 abundance. Our findings suggest 11q23.1 trans-eQTL targets comprise a *C11orf53-*related network that is likely tuft cell-specific and reduced expression of these genes correlates with reduced tuft cell abundance *in silico*.

## Introduction

Colorectal cancer (CRC) is the fourth most common cancer type UK and worldwide (Cancer Research UK, 2017; Rawla et al., 2019). Approximately 40% of CRC risk is attributable to heritable genetic variation (Graff et al., 2017), with rare, highly penetrant mutations responsible for a small fraction of overall risk (Jasperson et al., 2010; Sud et al., 2017). Genome wide association studies (GWAS) have identified 129 common genetic variants associated with CRC risk (Huyghe et al., 2019; Law et al., 2019). Several common CRC genetic risk variants are associated with heritable changes in the expression levels of genes in colonic mucosa, known as expression quantitative trait loci (eQTLs) (Closa et al., 2014; Hulur et al., 2015; Loo et al., 2017; Vaughan-Shaw et al., 2021).

Genetic variation at 11q23.1 is associated with increased CRC risk (Tenesa, Farrington et al., 2008). However, the large number of variants in high linkage disequilibrium at 11q23.1 makes identifying the causal variant, a key step to identifying the mechanism of gene dysregulation, difficult. Studies have shown that CRC risk associated variation at several 11q23.1 variants is correlated with the downregulation of three local genes; *C11orf53, COLCA1, COLCA2* (Biancolella et al., 2014; Loo et al., 2017; Smillie, 2015), known as cis-eQTL targets. We have recently shown that variation at rs3087967, a single nucleotide variant in the 3’UTR of *C11orf53*, is significantly correlated with distal CRC risk and a myriad of distant, trans-eQTL targets throughout the colon (Vaughan-Shaw *et al*., 2021), Additional File 1. Of these, only two have a common, well-described function; *IL17RB* and *TRPM5* are experimentally determined markers of tuft cells - a rare epithelial cell-type (Haber et al., 2017; Kaske et al., 2007). The function of several other trans-eQTL targets is currently unknown and their exact relevance to both 11q23.1 cis-eQTL target expression and CRC risk is not characterised.

Current methods of eQTL detection, while widely used, exhibit some key limitations. eQTLs are frequently identified by linear models of bulk RNA-seq/microarray transcriptome analysis of healthy tissue (Michaelson et al., 2009; Richards et al., 2012; Stranger et al., 2007). These methods often treat both gene expression and single-nucleotide polymorphisms as independent, linear entities: assumptions which over-simplify the complex relationships governing gene expression dynamics and limit results to additive gene-dosage related findings.

In addition, eQTL analyses require performance of an extremely large number of independent tests to be conducted, limiting its sensitivity. Correlation-based methods of gene expression analysis, such as weighted gene co-expression network analysis (WGCNA, Langfelder & Horvath, 2008), circumvent this problem by agnostically identifying correlations between individual genes, and entire, non-overlapping gene modules, with binarized categorical or quantitative traits. Modules of correlated genes may themselves be correlated with sample phenotypes. In addition, WGCNA also does not require a hard thresholding of correlations, a major advantage over other correlation-based methods that rely on arbitrary cut offs. We have recently shown that WGCNA can be efficacious at identifying genes driving transcriptional dynamic changes in the colorectum of patients treated with Vitamin D (Blackmur et al., 2022), even in the absence of statistically significant changes in differential expression analysis. Moreover, as CRC risk-associated eQTL targets are identified by bulk expression methods, findings are inherently limited in their potential to detect cell-specific changes, particularly in rare cell-types; expression changes within which may be masked due to low relative abundance.

We hypothesised that the expression of 11q23.1 trans-eQTL targets may be correlated with only a single cis-eQTL target, a relationship which may in turn be specific to a single epithelial cell-type in the colon. We utilised WGCNA across and within single-cell RNA sequencing (scRNAseq) clusters to further characterise eQTL target expression relatedness in colonic epithelial cell-types. An overview of the analysis performed in this study is shown in Figure 1.

**Figure 1.**
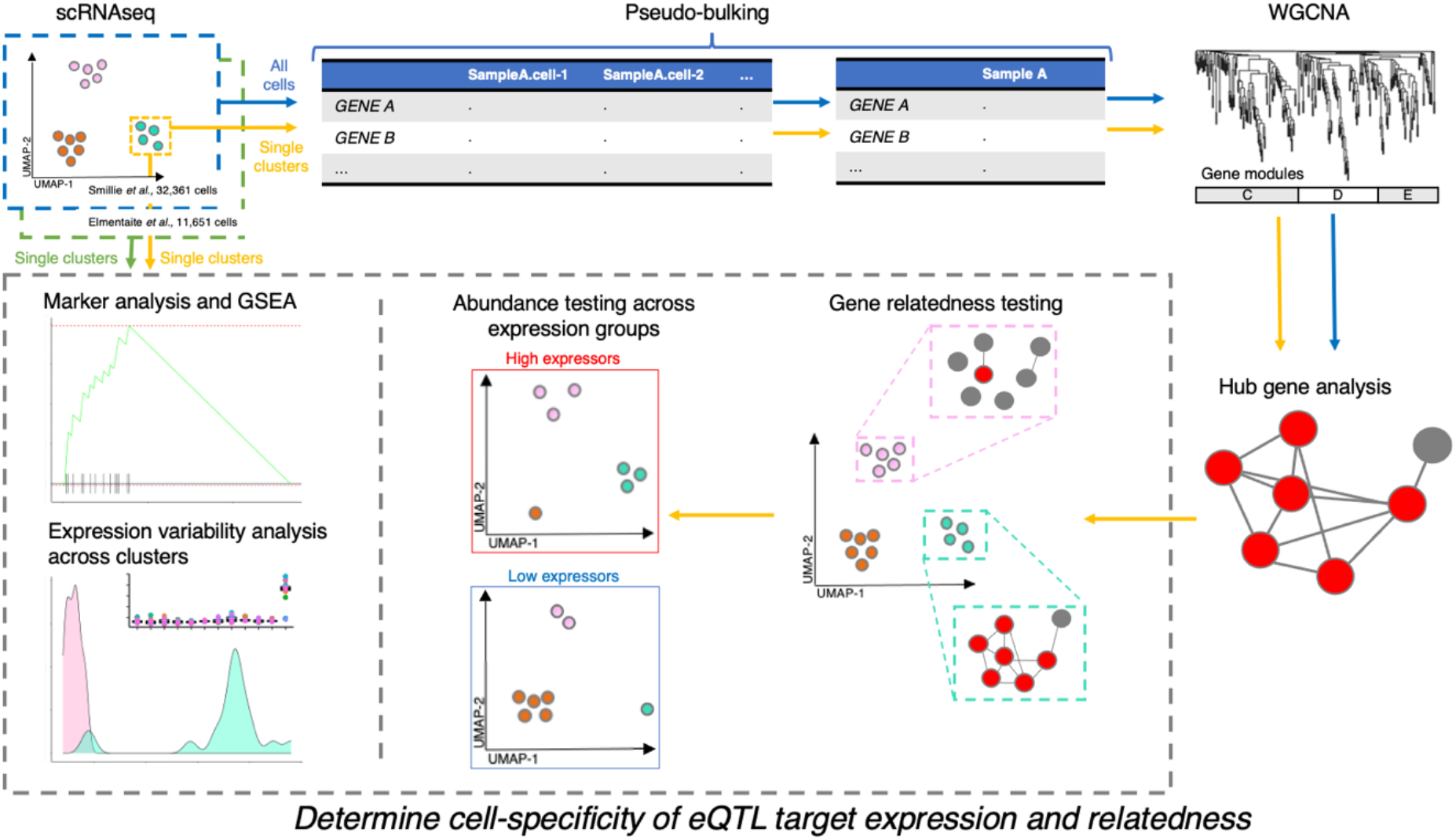
Overview of study design and analysis. This study makes use of two single cell RNA sequencing (scRNAseq) datasets: Smillie *et al*., (2019) – n=32,261 and Elmentaite *et al*., (2021) - n=11,651. The analysis performed on each dataset is outlined by arrow colour: blue=all cells from Smillie *et al*., (2019), yellow=individual clusters from Smillie *et al*., (2019), green=individual clusters from Elmentaite *et al*., (2021). WGCNA=Weighted Gene Co-expression Network Analysis, GSEA=Gene Set Enrichment Analysis.

## Results

### Understanding cis-eQTL specificity of 11q23.1 trans-eQTL effects

To test potential colonic epithelial cell-specificity of 11q23.1 eQTL effects, we sought to analyse the expression of target genes within colonic epithelial cell-types. To this end, we obtained scRNAseq from 32,361 healthy human colonic mucosa epithelial cells from eleven individuals (Smillie et al., 2019). To first assess the validity of this dataset in studying 11q23.1 variation-related expression dynamics, we devised a method of pseudo-bulking the expression from all cells in this dataset, aggregating the expression of every gene across all cells from each sample (see Figure 1 and Methods). Pseudo-bulked expression from 11q23.1 nominally significant trans-eQTL targets, present in the scRNAseq dataset (p<0.01, n=273), Figure 2A, was then subject to WGCNA (Langfelder & Horvath, 2008). Genes were agnostically grouped into modules of correlated expression, and the correlation between the eigenvector (first principal component) of each module with sample traits and pseudo-bulked cis-eQTL target gene expression was calculated, Figure 2B. The blue module, comprising 77 genes, was found to correlate with *C11orf53* expression (cor=0.81, FDR=2e-04), but none of the sample traits, *COLCA1* or *COLCA2* expression. The blue gene module includes 17 of the 20 significant 11q23.1 trans-eQTL targets that passed gene filtration quality control (FDR<0.05, Vaughan-Shaw *et al*., 2021); *ALOX5, SH2D6, TRPM5, BMX, PSTPIP2, GNG13, IL17RB, HTR3E, PTGS1, SH2D7, OGDHL, MATK, PLCG2, LRMP, PIK3CG, HTR3C* and *CAMP*, thus indicating the genes comprising this module are specifically related to the expression of *C11orf53*, Supplementary Table 2.

**Figure 2.**
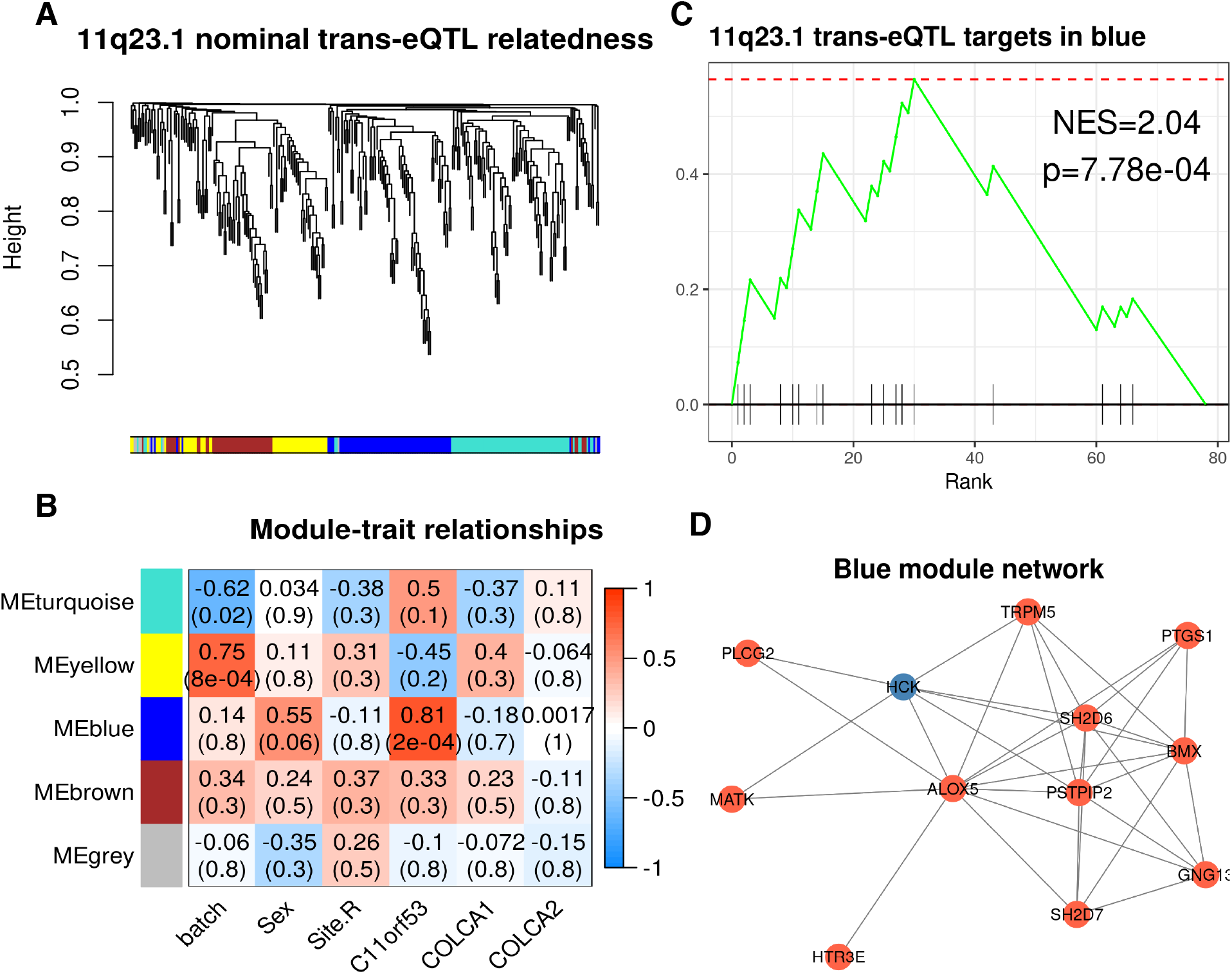
11q23.1 trans-eQTL targets are correlated with the expression of *C11orf53*, but not *COLCA1* or *COLCA2* in pseudo-bulked scRNASeq. (A) Hierachical clustering of complete distances between pairwise correlations of 11q23.1 nominal trans-eQTL targets pseudo-bulked expression (p<0.01, n=273) in 32,361 healthy colonic epithelial scRNAseq. (B) Weighted Gene Co-expression Network Analysis (WGCNA) identified module trait relationships. Pearson correlations shown above Benjamini-Hochberg corrected p-values in brackets. (C) Gene Set Enrichment Analysis (GSEA) of 11q23.1 trans-eQTL targets (FDR<0.05) in blue module genes, ranked by their module membership. (D) Kamada-kawai network of Blue module relatedness (adjacency > 0.3). Red nodes indicate FDR<0.05 trans-eQTL target of 11q23.1.

To assess the contribution of 11q23.1 genes to the correlation of the blue module with *C11orf53*, we performed gene set enrichment analysis of the 11q23.1 trans-eQTL target genes (FDR<0.05) against the genes in the blue module, ranked by their module membership - a measure of their relatedness to all other genes in the module (Langfelder & Horvath, 2008). We found the genes in this module are highly enriched for 11q23.1 trans-eQTL targets, normalised enrichment score (NES) = 2.04, p=7.78e-04, Figure 2C. In addition, 11 trans-eQTL targets comprised the 12 genes with the greatest adjacency in the blue module (adjacency > 0.3), Figure 2D. We also find that module membership is highly correlated with gene significance for *C11orf53* in the blue module, reinforcing the overall correlation of the network with *C11orf53* expression (Supplementary Figure 1). Together, this replicates the trans-eQTL target expression relatedness previously described (Vaughan-Shaw et al., 2021) and indicates that the expression of the majority of significant 11q23.1 trans-eQTL target genes is correlated with *C11orf53* specifically in this dataset.

### Analysing cell-specificity of 11q23.1 eQTL target expression

To test the potential for cell-specific expression of 11q23.1 eQTL targets in this dataset, we performed dimensionality reduction and clustering of the single cells, identifying a total of 12 transcriptionally distinct cell-clusters, named ‘0’-’11’, Figure 3A. We then calculated the markers of each cluster and found the markers of cluster 11, comprising 318 cells, includes fourteen 11q23.1 trans-eQTL targets (FDR<0.05), Table 1 (full list of cluster 11 marker genes available in Additional File 2). In addition, cluster 11 markers are significantly enriched for 11q23.1 trans-eQTL targets (NES=2.15, p=7.52e-06), Figure 3B. Cluster 11 was found to transcriptionally resemble the Smillie *et al*., tuft cell cluster, by enrichment of each cluster’s markers with the authors’ putative markers (NES=2.37, FDR=1.23e-06, see Methods). Together, this indicates cluster 11 is transcriptionally defined by the expression of 11q23.1 trans-eQTL targets, several of which are themselves putative tuft cell markers.

**Figure 3.**
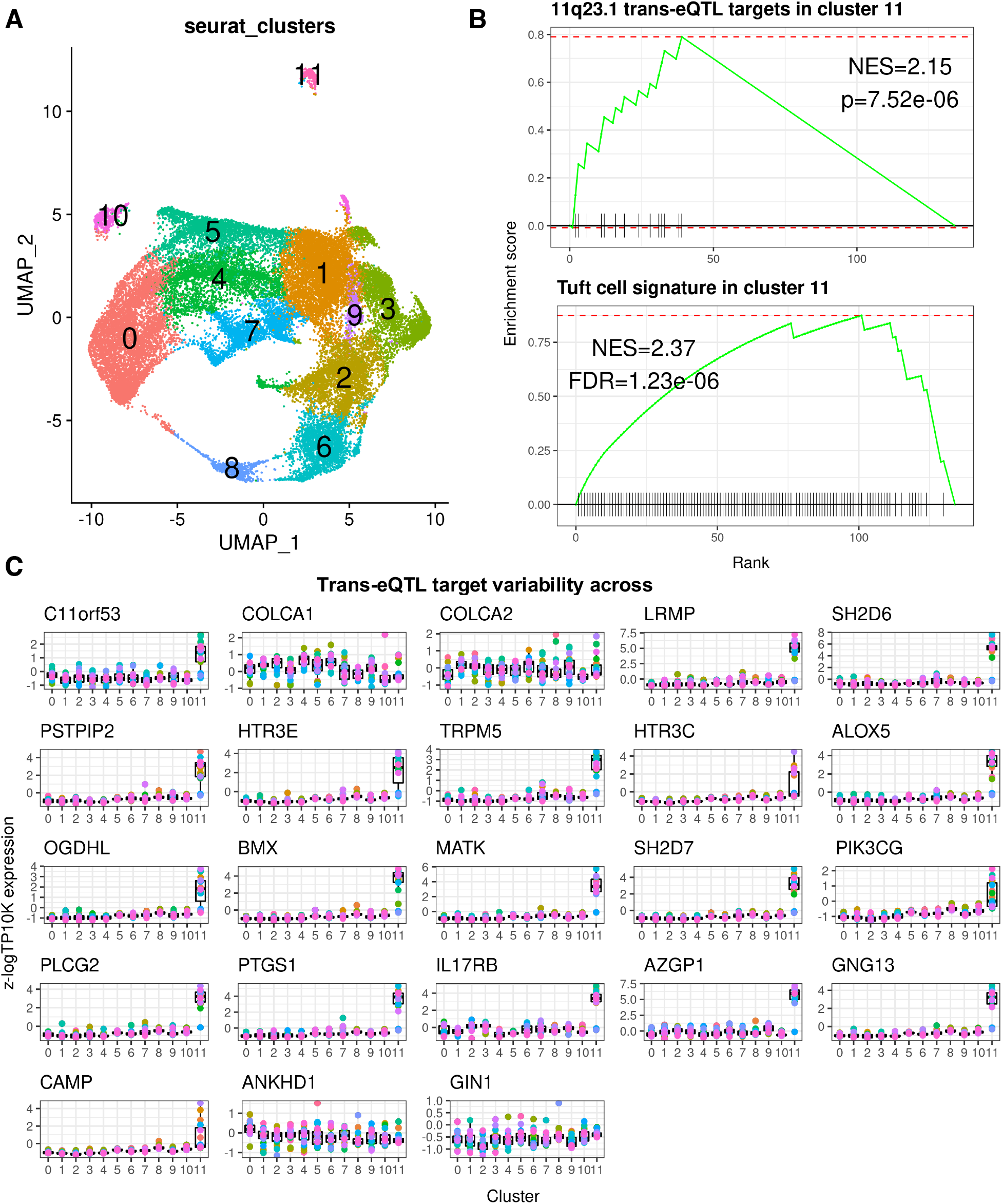
11q23.1 trans-eQTL expression demarcates tuft-like cell cluster. (A) UMAP of 32,361 epithelial scRNASeq, coloured by cell cluster - identified using Seurat (Hao et al., 2021). (B) Upper: GSEA of 11q23.1 trans-eQTL targets (Vaughan-Shaw *et al*., 2021; FDR<0.05) in cluster 11 markers. Lower: GSEA of putative colonic tuft cell signature (Smillie *et al*., 2019) in cluster 11 markers. P=P value, FDR=False Discovery Rate (C) Relative, pseudo-bulked expression (log transcripts per 10,000) of 11q23.1 trans-eQTL targets (FDR<0.05) across clusters.

**Table 1.** 11q23.1 trans-eQTL targets identified as cluster 11 markers. Markers calculated using MAST (McDavid *et al*., 2020). Avg_log2FC=average log2 fold change between cluster 11 and all other clusters. Pct1=Proportion of gene expression in cluster 11. Pct2=Proportion of expression in non-cluster 11. P_val_adj=FDR-corrected p-value.

| Gene | p_val | avg_log2FC | pct.1 | pct.2 | p_val_adj |
| --- | --- | --- | --- | --- | --- |
| *AZGP1* | 1.00E-279 | 2.5735445 | 0.72 | 0.029 | 1.00E-279 |
| *SH2D6* | 1.00E-279 | 2.43533298 | 0.736 | 0.007 | 1.00E-279 |
| *LRMP* | 1.00E-279 | 1.97284472 | 0.572 | 0.005 | 1.00E-279 |
| *PTGS1* | 1.18E-278 | 1.33766533 | 0.459 | 0.003 | 1.75E-274 |
| *IL17RB* | 1.00E-264 | 1.04046297 | 0.412 | 0.031 | 1.48E-260 |
| *BMX* | 7.18E-248 | 1.18575976 | 0.406 | 0.002 | 1.07E-243 |
| *ALOX5* | 8.25E-244 | 1.09872574 | 0.409 | 0.003 | 1.22E-239 |
| *MATK* | 2.88E-240 | 1.3710223 | 0.406 | 0.003 | 4.28E-236 |
| *SH2D7* | 2.50E-235 | 1.20590753 | 0.393 | 0.002 | 3.72E-231 |
| *GNG13* | 2.93E-220 | 0.98868335 | 0.362 | 0.001 | 4.34E-216 |
| *TRPM5* | 1.32E-214 | 0.94972732 | 0.377 | 0.004 | 1.96E-210 |
| *PLCG2* | 1.22E-209 | 0.98215989 | 0.365 | 0.002 | 1.82E-205 |
| *PSTPIP2* | 2.21E-197 | 0.87613246 | 0.333 | 0.004 | 3.28E-193 |
| *HTR3E* | 4.86E-184 | 0.8684383 | 0.308 | 0.001 | 7.21E-180 |

Because the expression of 11q23.1 trans-eQTL targets is reduced in individuals with the CRC risk-associated genotype at 11q23.1 (Vaughan-Shaw et al., 2021), we wanted to assess the variability in the expression of these genes within each cluster. We pseudo-bulked the expression of all cells from each sample, within clusters, and analysed 11q23.1 trans-eQTL target expression (Figure 3C). The relative expression level and variability of *C11orf53* and 18 of the 11q23.1 trans-eQTL targets is overwhelmingly greatest within cluster 11, indicating the eQTL effect on these genes is exacerbated within this cluster. Notably, the relative variation and expression of cis-eQTL targets *COLCA1* and *COLCA2* and trans-eQTL targets *ANKHD1* and *GIN1*, is not greatest within this cluster, suggesting the eQTL effect on these genes may not be driven by transcriptional dynamics within this cell-type. In addition, we analysed 11q23.1 eQTL target variability at the single cell level, Supplementary Figure 2. The variability of *C11orf53* and the same 18 trans-eQTL targets was found to be greatest within cluster 11, replicating the potential exacerbation of the eQTL effect within this cluster and supporting the validity of our pseudo-bulk method. *C11orf53* expression, however, was not found to demarcate cluster 11 by marker identification analysis (see Methods), data not shown.

To test the robustness of tuft cell-like mapping of 11q23.1 trans-eQTL target expression and variability, we replicated this analysis in an independent dataset of 11,651 healthy adult colonic epithelial cells from 3 individuals (Elmentaite et al., 2021), Supplementary Figure 4A. In this instance, we identified 19 cell-clusters by dimensionality reduction and unsupervised clustering, named ‘0’-’18’. The markers of cluster 18 were significantly enriched for the expression of 11q23.1 trans-eQTL targets (NES=2.50, P=5.52e-09, Supplementary Figure 4B), 11 of which were identified as markers of this cluster, Supplementary Table 3. Cluster 18 was also enriched for the tuft cell signature (NES=2.41, P=7e-08), replicating our previous finding. The relative variability and expression of the majority of 11q23.1 trans-eQTL targets was also greatest within cluster 18, Supplementary Figure 4C. The expression of 13 trans-eQTL targets was also most variable within cluster 18 when expression was analysed at the single cell level, Supplementary Figure 4. Interestingly, *C11orf53* expression variability was not found to vary greatest within cluster 18 when using expression from single cells.

### Understanding 11q23.1 cis- and trans-eQTL relatedness within clusters

The demarcation and variability of 11q23.1 trans-eQTL target expression within a tuft-like cell cluster strongly indicates the eQTL effect is derived from altered gene expression in this cell-type specifically. However, it is possible that the gene-gene relatedness of 11q23.1 eQTL targets, including those with *C11orf53*, is not specific, but rather exacerbated in this cluster. To test this, we sought to divide samples by genotype at rs3087967, the variant associated with expression changes of trans-eQTL targets (Vaughan-Shaw *et al*., 2021), and analyse the congruence of trans-eQTL target expression relatedness within clusters. While genotype information was unavailable for the Smillie *et al*., (2019) dataset, raw sequencing reads were available for the Elmentaite *et al*., (2021) dataset and as rs3087967 lies within the 3’ UTR of *C11orf53*, we performed variant calling on these samples using orthogonal tools, Supplementary Table 4. Using freebayes (Garrison & Marth, 2012), we found that all samples were called as homozygous for the non-risk allele at rs3087967, except for one sample called as heterozygous. However, as all 3 other samples from this individual were called as homozygous non-risk, it is likely this is a technical error. Using bcftools (Danecek et al., 2021), all samples were identified to be homozygous non-risk for rs3087967. The lack of genetic variation at rs3087967 in these samples is consistent with the absence of heightened *C11orf53* variability in cluster 18 and indicates this dataset would be unlikely to be of use to identify *C11orf53-*related expression dynamics. The relatively high variability of trans-eQTL target expression within cluster 18 may be indicative of non-11q23.1-related dynamics, such as changes during differentiation or cell cycle progression.

To test the potential efficacy of the Smillie *et al*., (2019) dataset to further study the 11q23.1 eQTL target expression dynamics, we compared the standardized variability of 11q23.1 trans-eQTL targets within the respective demarcated clusters at the single-cell level, Supplementary Table 5. We found that the expression variability of 14 of the 15 11q23.1 trans-eQTL targets, expressed in both Elmentaite *et al*., (2021) cluster 18 and Smillie *et al*., (2019) cluster 11, was substantially increased in the latter (fold-change range 1.46-9.7, median=1.97, mean=2.83, 95% confidence interval=1.50-4.18). The only 11q23.1 trans-eQTL target not to exhibit increased variability in Smillie *et al*., (2019) cluster 11 was OGDHL (fold change=0.73). Notably, the expression of *C11orf53*, was also greater in Smillie *et al*., cluster 11 (fold change=1.94). Subsequent analysis was therefore focussed on the Smillie *et al*., (2019) dataset.

To divide samples into high and low expressors of *C11orf53*-expression related genes, we hierarchically clustered samples based on the relative expression of pseudo-bulk WGCNA-identified blue module hub genes. Hub genes were defined by module membership > 0.7, intramodular connectivity > 0.7 and network adjacency > 0.3 and include 10 genes: *TRPM5, PSTPIP2, SH2D6, ALOX5, BMX, GNG13, SH2D7, HCK, PLCG2, MATK*, Figure 4A. Five samples were separated at the first branch of clustering and exhibit a strong relative reduction in the expression of these genes. This grouping of samples is henceforth referred to as the ‘blue module hub gene grouping’. To assess the significance of this separation at representing underlying transcriptional differences, we performed 10,000 permutations, using 10 randomly sampled genes, and generated a p-value of 0.055.

**Figure 4.**
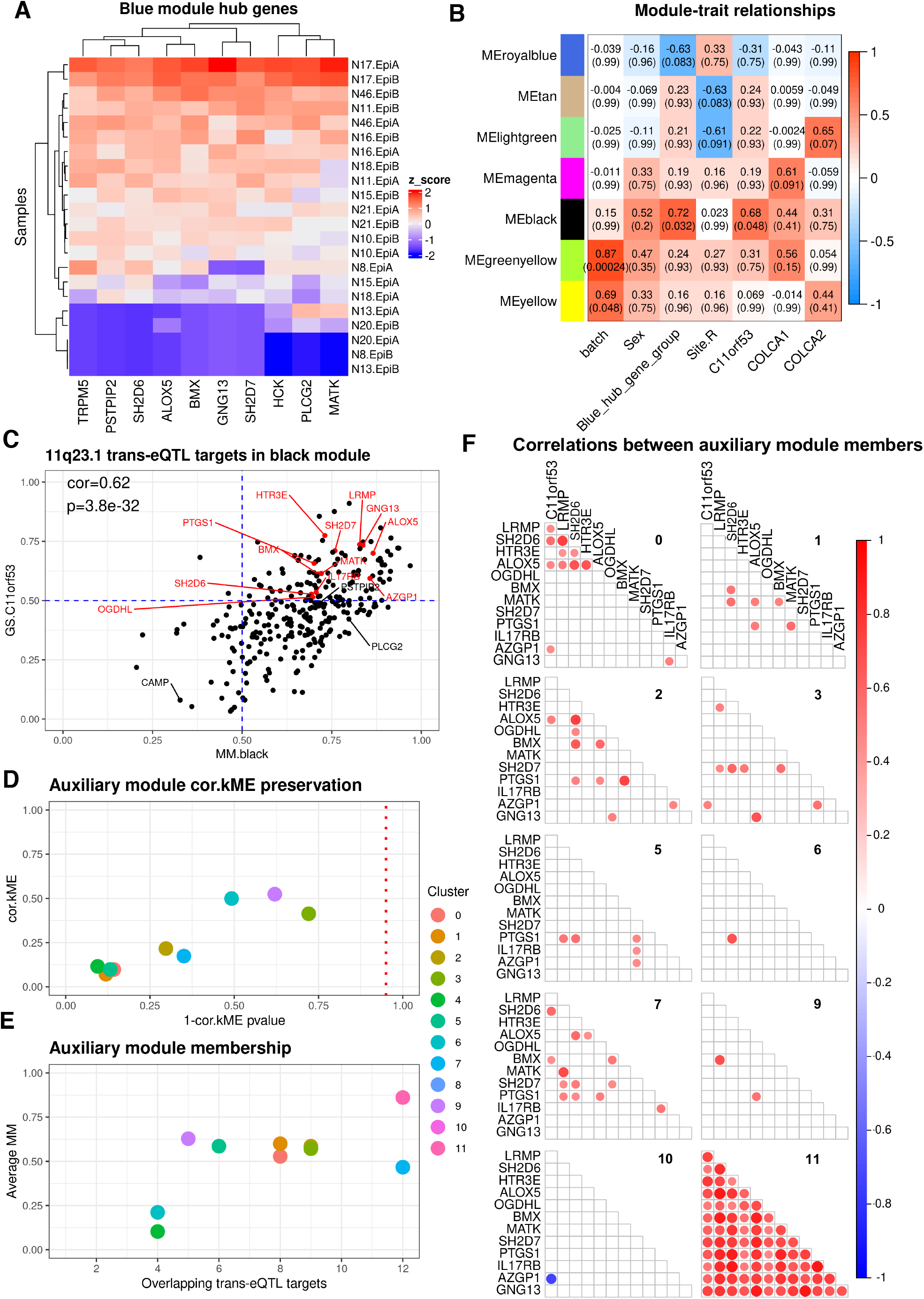
Several 11q23.1 trans-eQTL targets comprise *C11orf53-*correlated network in tuft-like cluster only. (A) Relative (z-score) expression of blue module hub genes from Fig.1D (MM>0.7, kIM>0.7, adj > 0.3). Robustness assessed by 10,000 permutations: p=0.055. (B) Module trait matrix of gene modules identified by WGCNA within pseudo-bulked expression from cluster 11. Only modules exhibiting a correlation with a covariate (FDR<0.1) are shown. Total number of modules=20. (C) Correlation of Gene Significance (GS) and Module Membership (MM) of genes in the black module from (B). 11q23.1 trans-eQTL targets are highlighted. 11q23.1 trans-eQTL targets with GS.C11orf53 > 0.5 and MM.black > 0.5 (red) were used as the auxiliary module. (D) Preservation of correlations between auxiliary module genes with the module eigengene (ME) and equivalent values in cluster 11. Dashed line indicates nominal significance threshold (p=0.05). (E) Average MM of genes within the auxiliary module across clusters. (F) Pairwise pearson correlations (p<0.05) between the pseudo-bulked expression of auxiliary module genes across clusters. Cluster number in upper right of each facet. Cluster 4 and 8 exhibited no significant correlations and so are not plotted.

To first test gene-gene relatedness within cluster 11, we performed WGCNA on the relative, pseudo-bulked expression of the top 5000 most variable genes within this cluster, identifying a total of 20 modules, 7 of which exhibited correlations with sample covariates that approached significance (FDR<0.1), Figure 4B. We found one module, ‘cluster 11 black’, which was highly correlated with both the ‘blue hub gene grouping’ (cor=0.72, FDR=0.032) and *C11orf53* expression (cor=0.68, FDR=0.048). ‘Cluster 11 black’ is comprised of 290 genes, including fifteen 11q23.1 trans-eQTL targets, Additional File 3. As the ‘blue module hub gene’ grouping is derived from analysis of genes correlated with *C11orf53* across all cells, the correlation of ‘cluster 11 black’ with both this grouping and *C11orf53* expression indicates that this relationship is preserved within and potentially derived from this cell-cluster.

We then wanted to test whether the gene-gene relatedness of 11q23.1 trans-eQTL targets, correlated with *C11orf53*, was specific to cluster 11. To this end, we defined an auxiliary module, comprised of the twelve 11q23.1 trans-eQTL targets correlated with *C11orf53* (cor>0.5, p<0.05) in cluster 11 black, which exhibited high module membership (MM.black > 0.5), Figure 4C. These genes include: *HTR3E, LRMP, GNG13, ALOX5, SH2D7, PTGS1, MATK, BMX, AZGP1, IL17RB, SH2D6* and *OGDHL*. We performed pairwise module preservation of this auxiliary module, using identical parameters as used for intra-cluster 11 analysis, in all other clusters, see Methods. For two modules, cluster 8 and cluster 10, the 5000 most variable genes included only a single member of the auxiliary module and so were excluded from this analysis. To analyse the preservation of the connectivity of these genes, we assessed the similarity of each gene correlation with the module eigengene, and the equivalent values in cluster 11 (cor.kME), Figure 4D. No module exhibited a significant cor.kME with cluster 11 (p>0.05), indicating there is low overall preservation of the connectivity between these genes in all other clusters. The average module membership of genes in this module (average.MM) was also reduced in all clusters compared with cluster 11, Figure 4E. Finally, we analysed the pairwise gene-gene correlations between all members of the auxiliary module and *C11orf53* within each cluster, Figure 4F. While there are rare correlations between these genes in other modules (cor>0.5, p<0.05), all comparisons reach this threshold in cluster 11. Taken together, this evidence indicates these 12 trans-eQTL targets comprise a transcriptional network that is correlated with *C11orf53* expression and likely specific to cluster 11.

### Identification of cluster 11 abundance associated genes

As many of the 11q23.1 eQTL targets, including those comprising the cluster 11-specific network, demarcate this cluster, we wanted to examine the relationship between their expression and cluster 11 abundance. First, we performed linear modelling of the relative abundance of cluster 11 and the pseudo-bulked expression of *C11orf53* only. We found that the expression of *C11orf53* is associated with the relative abundance of cluster 11 (coefficient=0.389, p=0.00431) indicating potential for the expression of *C11orf53* alone to be moderately predictive of the abundance of this cell-type, Figure 5A.

**Figure 5.**
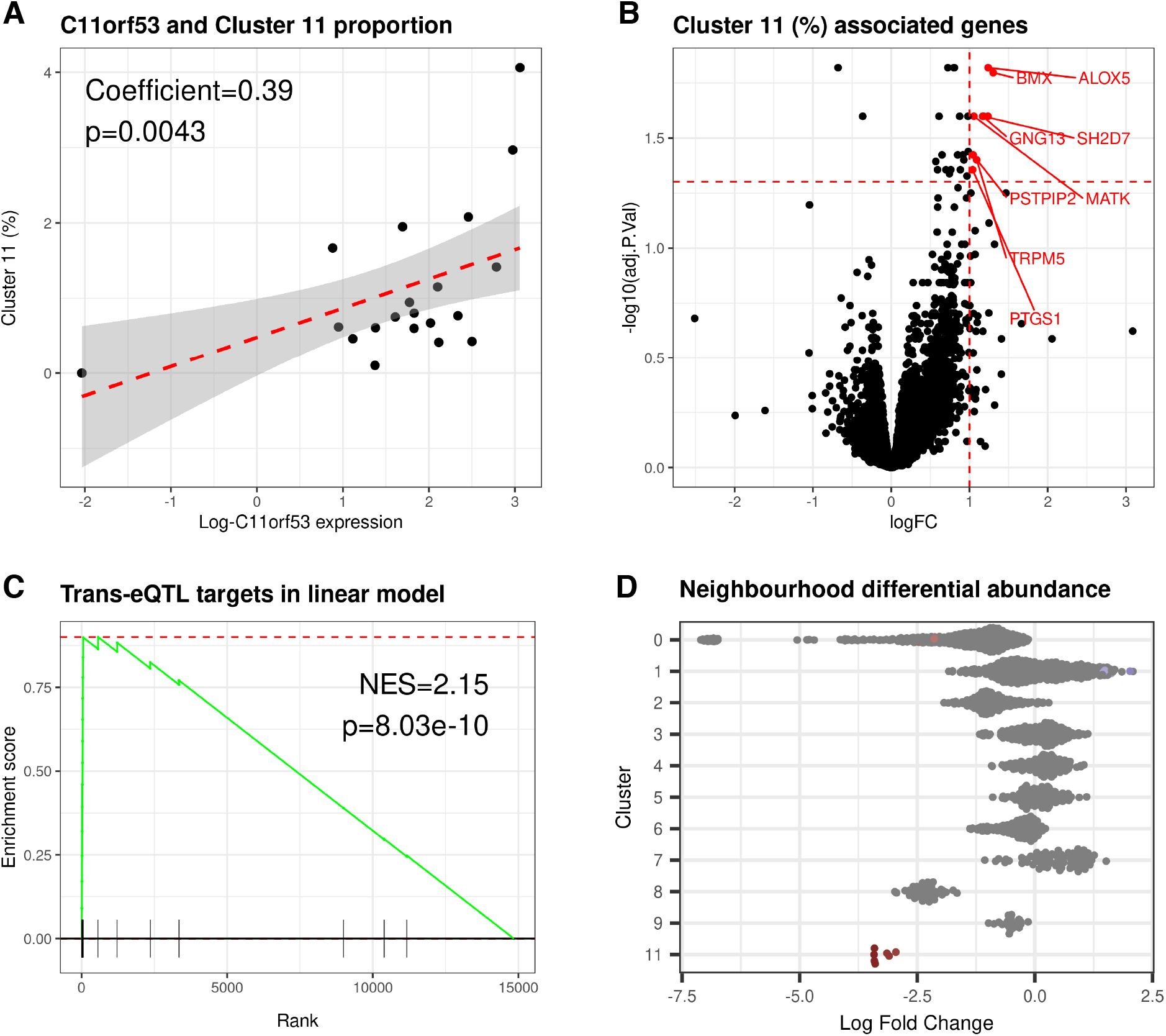
11q23.1 trans-eQTL target expression is associated with the abundance of tuft cell-like cluster. (A) Linear modelling of pseudo-bulked *C11ORF53* expression and cluster 11 abundance. (B) Volcano plot of linear model result on pseudo-bulked expression of all 14,843 genes. Significantly associated 11q23.1 trans-eQTLs (logFC>1, FDR<0.05) highlighted. (C) GSEA of 11q23.1 trans-eQTLs (FDR<0.05) in (B), ranked by logFC. (D) Differential abundance of neighbourhoods in the ‘Low’ blue hub gene group compared to ‘High’. Neighbourhoods were identified using miloR (Dann *et al*., 2021) and grouped by cluster. Only significant neighbourhoods with major cluster proportion > 0.8 are plotted and only significant neighbourhoods (Spatial FDR < 0.01) are coloured by their logFC (red=down, blue=up). No neighbourhoods from cluster 10 were comprised of major proportion > 0.8.

To agnostically test the predictive power of 11q23.1 trans-eQTL target expression on cluster 11 abundance, we tested the association between the expression of all genes and the proportion of cluster 11 in samples, Figure 5B. We found that all genes significantly associated with the abundance of this cluster (FDR<0.05, log-fold change>1) were indeed 11q23.1 trans-eQTL targets, including: *ALOX5, BMX, GNG13, MATK, SH2D7, PSTPIP2, TRPM5* and *PTGS1*. In fact, the strength of gene association with cluster 11 abundance was also significantly enriched for 11q23.1 trans-eQTL targets (NES=2.15, p=8.03e-10, Figure 5C).

While our linear modelling strongly supports a predictive role of 11q23.1 eQTL target expression in the abundance of cluster 11, we wanted to agnostically test whether the expression of the *C11orf53*-related trans-eQTL targets was correlated with abundance changes to any cluster. To this end, we utilised miloR (Dann et al., 2021) to calculate cell-neighbourhoods, Supplementary Figure 5, which were then used to perform differential abundance testing across the ‘blue module hub gene grouping’, defined in Figure 4A. To generalise neighbourhoods to the cell clusters we have identified, we subsequently filtered for neighbourhoods which represented a majority of a single cluster (majority proportion > 0.8) Figure 5D. No neighbourhood from cluster 10 was comprised of a majority proportion of 0.8 and so this cluster was not included. In the low *C11orf53*-related gene expressing samples there are significant reductions (Spatial FDR < 0.01) from cluster 0 and increases in neighbourhoods in clusters 1, Figure 5D. The change in abundance of these neighbourhoods represent a small proportion of the total number of neighbourhoods detected in these clusters (2.1% and 1.3% respectively) and are therefore unlikely to represent a dramatic phenotype.

In contrast, all 7 of the neighbourhoods comprising a majority of cluster 11 cells were significantly underrepresented in the low ‘blue module hub gene group’. These results indicate 11q23.1 eQTL target expression is correlated with a significant and likely specific change in abundance of cluster 11 cells.

## Discussion

In this study, our pan-cluster WGCNA serves as validation of the expression relatedness between trans-eQTL targets previously identified to be correlated with CRC associated variation at 11q23.1 (Vaughan-Shaw *et al*., 2021). 11q23.1 trans-eQTL target expression was also found to be more correlated with *C11orf53* over other cis-eQTL targets. Following clustering of single cells, many of the genes demarcating a single cluster, number 11, were found to be 11q23.1 trans-eQTLs targets. Enrichment of these markers against putative gene sets showed cluster 11 transcriptionally resembled tuft cells, replicated in an independent dataset. WGCNA within cluster 11 identified several 11q23.1 trans-eQTL targets that exhibited a high level of relatedness, subsequent analysis of the preservation of this relatedness indicated this was likely to be specific to this cell cluster. Finally, samples which exhibited low overall expression of 11q23.1 trans-eQTL targets most correlated with one another, were found to exhibit a specific and dramatic reduction of the tuft cell-like cluster. Therefore, our results potentiate CRC risk associated variation at 11q23.1 reduces the abundance of colonic tuft cells.

To our knowledge, this is the first study to map CRC risk associated eQTL targets to specific epithelial cell-types. Heritable inflammatory bowel disease risk loci have recently been associated with shifts in transcriptional dynamics in individual colonic epithelial cells (Smillie et al., 2019) and it is possible that other CRC risk variants, with robust eQTL effects, are associated with cell-specific changes in transcriptional dynamics. Delineating the cell-specific expression of CRC risk-associated eQTLs may provide valuable insights into the mechanism of risk-associated pathophysiology and should be a focus of future work. The growing size and availability of scRNAseq datasets will likely make cell-specificity mapping of heritable disease associated eQTLs significantly easier and available, particularly if genotype data is available. Indeed, our study is also not the first use of WGCNA to detect gene-gene relatedness in scRNAseq data. WGCNA has been utilised to identify gene modules associated with the activation of neuronal stem cells (Luo et al., 2015) and human induced pluripotent stem cells (Wu et al., 2021), however, like our own study, these studies did not utilise expression from individual cells as the input for WGCNA.

Surprisingly, the expression of the vast majority of 11q23.1 eQTL targets mapped to the cell-type that transcriptionally resembles tuft cells, including; *LRMP, IL17RB, SH2D6, PLCG2, PSTPIP2, TRPM5, SH2D7, AXGP1, PTGS1, ALOX5, BMX*. Many of the most significant 11q23.1 trans-eQTL targets, such as *LRMP, SH2D7* and *ALOX5*, have not previously been associated with specific expression in this cell-type, potentiating their status as a marker in the colonic epithelium. Additional markers of the tuft cell-like cluster include *HCK* and *HPGDS*, for which there is some orthogonal evidence of expression within tuft cells (Gerbe et al., 2011; Yamaga et al., 2018). This improves our confidence that cluster 11 represents this cell-type and is not an artefact of the analysis.

It is noteworthy that many of the genes with expression found to be in association with *C11orf53* in the scRNAseq data were indeed identified as trans-eQTL targets in the bulk analysis (Vaughan-Shaw et al., 2021). While the mapping of the expression of these genes to a tuft-like cell cluster could only be achieved by using expression from individual cells, the prior identification of these genes by bulk analysis is a testament to the power of such approaches, and concordance of their findings with single cell-based methods.

Finally, the overall potentiation of tuft cell perturbation is of great interest regarding characterisation of the mechanism governing CRC risk at 11q23.1. Tuft cells are associated with stem-, neuro-transmitting and immune-related functions (Banerjee et al., 2020; Gerbe et al., 2016; Middelhoff et al., 2017; Yi et al., 2019), however much of the evidence regarding their function is derived from other organs and cannot necessarily be extrapolated to the colon. Interestingly, genetic ablation of tuft cell abundance has been associated with exacerbated tumour progression in mouse models of pancreatic cancer (DelGiorno et al., 2020; Hoffman et al., 2021). Both studies show this is likely in association with perturbed immune cell function and signalling. In line with this, tuft cell abundance has recently been shown to be reduced in quiescent ulcerative colitis patients (Kjærgaard et al., 2021), suggesting tuft cells are involved in immune regulation in the colon. Future work should aim to experimentally validate the relationship between 11q23.1 variation and tuft cell abundance, examine how this impacts tumourigenesis and identify the potential mechanism of CRC risk predisposition.

## Materials and methods

### Availability of data and materials

Raw gene-level expression counts of healthy colonic epithelial cells, published by Smillie *et al*., (2019), were obtained from Single Cell Portal using accession code SCP259. Samples from one patient, N51, were removed as they were found to be outliers on the basis of cell-level mitochondrial and ribosomal protein gene expression, in addition to principal component analysis of pseudo-bulked expression. Fastq files for the second dataset, published by Elmentaite *et al*., (2021) were obtained from https://www.ebi.ac.uk/arrayexpress, using accession code E-MTAB-9543. Fastq files were aligned to the hg19 transcriptome using the 10X Genomics Cell Ranger v3.02 pipeline (Zheng et al., 2017) to produce the raw gene-level counts. All code used for the analysis performed in this study is available at https://github.com/BradleyH017/Harris_et_al_WGCNAscRNA.

### Pre-processing, dimensionality reduction and clustering of scRNAseq data

All subsequent expression analysis was completed in R version 4.0.2 (R Core Team, 2020). Once raw counts had been obtained for both datasets, bad quality droplets were filtered by a series of quality control steps: i) potential empty droplets were detected by thresholding at the inflection point of the barcode rank plot of cells, calculated using *DropletUtils* v1.8.0 (Griffiths et al., 2018), ii) genes expressed in fewer than 20 cells were removed, iii) cells with an expression sparsity > 0.99 were removed, iv) cells with proportion of mitochondrial gene expression greater than 2.5*(median absolute deviation) from the median proportion were removed.

Following filtration, counts were loaded into a Seurat object using Seurat v4.0.1 (Hao et al., 2021). Initial clustering was performed using Seurat’s reciprocal PCA method of batch correction with *SCTransform* as per Hao *et al*., (2021) guidelines (https://satijalab.org/seurat/articles/integration_rpca.html). The processed Seurat object was first split by sample and data integration was performed on the first 50 principal components. The principal components of the integrated data were then calculated, which was used to calculate the UMAP embedding for all cells, based on 50 principal components

To identify clusters, a nearest neighbour graph was constructed on the integrated data, using 50 principal components. Clusters were then identified via the *FindClusters* function using a resolution of 0.6. For the Smillie *et al*., (2019) data analysis, we utilised a k value of 250, as this is concordant with the authors analysis and provided the greatest confidence in identification of clusters than other k values. Concordance was tested by enrichment against the authors cluster markers; data not shown. For the Elmentaite *et al*., (2021) analysis, we utilised a k value of 20. Any value greater than this resulted in failure to detect the tuft-cell resembling cluster.

To identify potential doublets in the filtered dataset, we utilised *DoubletFinder* v2.0.3 (McGinnis et al., 2019) as per the authors guidelines (https://github.com/chris-mcginnis-ucsf/DoubletFinder). The fully filtered dataset was then re-used as input for integration, dimensionality reduction and clustering as described above.

To test the robustness of the clusters we identified, we performed pairwise differential gene expression analysis by a receptor operator curve test using Seurat’s *FindMarkers* function. In order to only merge clusters which were extremely similar, without over-clustering, we defined similar clusters as those with less than 30 differentially expressed genes with area under the curve score of 0.6 differentiating them. We found no clusters that fall below this threshold and so did not modify our initial clustering in either dataset.

### Pan-cluster WGCNA

To analyse correlation of gene expression across all cells from the filtered Smillie *et al*., (2019) data, we first subset for the nominally significant 11q23.1 trans-eQTLs previously reported (Vaughan-Shaw et al., 2021). The relative pseudo-bulked expression was then calculated by: i) summing reads for each gene across all cells within samples, ii) recombining the summed reads into a non-normalised bulk matrix, iii) log normalising using TMM normalised size factors, calculated using *edgeR* v3.32.1 (McCarthy et al., 2012). Log-TMM normalised bulk expression was then z-scored across genes before analysis.

To perform the network analysis, we used *WGCNA* v1.69 (Langfelder & Horvath, 2008). First, *C11orf53, COLCA1* and *COLCA2* pseudo-bulk expression was extracted. A soft threshold of 14 was then selected after computation of mean connectivity and scale free topology, using the recommended ‘powerEstimate’. A signed adjacency matrix was then calculated, which was subsequently used to calculate a topological overlap matrix (TOM). Modules were defined by using a dynamic tree cut of hierarchically clustered gene expression by average distances. We did not find any modules with a height of module separation below 0.25 and so did not merge any module. Module eigengenes were then calculated and their correlation with binarized sex, batch, site, and relative, pseudo-bulked expression of *C11orf53, COLCA1* and *COLCA2* was then assessed. Correlation p-values were multiple testing corrected by the Benjamini-Hochberg method (Benjamini & Hochberg, 1995). To visualise the blue module hub genes, a network object was generated from the TOM of blue module genes using *network* v1.17.1 (Butts, 2008). Non-connected genes, along with adjacencies < 0.3, were then removed and remaining genes were plotted using *ggplot2* v3.3.5 (Wickham, 2009).

### Gene set enrichment analysis

All gene set enrichment analysis was carried out using R package fgsea v1.14.0 (Korotkevich et al., 2021). Genes were ranked by their module membership, gene significance for *C11orf53* expression, or log fold change of differential expression as stated. In the case where multiple gene sets were being tested, i.e cluster 11 markers against all Smillie et al., putative markers, p-values were multiple testing corrected by the false discovery rate method.

### Calculation of cluster markers

To calculate the markers of each of our own and the Smillie *et al*., (2019) clusters, we first generated a log-transcripts per 10,000 expression matrix for the expression of genes within each cell. This was done so that markers could be calculated using expression values that were not affected by the relative expression of genes within this dataset and so were more applicable to future use. Markers were identified by differential expression of genes within each cluster and all other clusters combined, using MAST v1.160 (Finak et al., 2015).

### Analysing the variability of trans-eQTL targets within clusters

To analyse the variability of the expression of 11q23.1 trans-eQTL targets within each cluster, we applied the pseudo-bulking approach described in *Pseudo-bulk WGCNA* to each cluster independently. To make expression variability comparable across samples and clusters, the expression was z-scored across samples.

To analyse expression variability at the single level, we utilised Seurat’s *FindVariableFeatures* function and variance stabilising transformation. In concordance with the identification of many trans-eQTL targets identified as markers, their mean expression was very low in several clusters within each dataset. For within dataset comparison of variance, we therefore utilised the raw variance, not normalising for mean expression. For the between dataset comparison of variance, we utilised the normalised variance values.

### *Genotyping of* *Elmentaite et al.,* (2021) *samples*

To identify the rs3087967 genotype of Elmentaite *et al*., (2021) samples, we utilised two variant calling approaches. These were chosen on the basis of findings from a recent review which identified these methods as the most sensitive for this purpose (Liu et al., 2019). Freebayes (Garrison & Marth, 2012) was used with default settings, genotyping over a 10bp region including rs3087967. Bcftools variant calling (Danecek et al., 2021) was performed on chromosome 11 using a minimum base quality of 30, disabling read-pair overlap detection and not discarding anomalous pairs.

### Sample group definition

In the absence of genotype data for Smillie *et al*., (2019) patients, we grouped samples into high and low expressors of a *C11orf53-*associated signature by defining the hub genes of the pseudo-bulk WGCNA blue module. These were defined by a module membership (MM) > 0.7, intramodular connectivity (kIM) > 0.7 and network adjacency > 0.3. Samples were then hierarchically clustered by their relative pseudo-bulked expression of these genes using complete distance. We tested the robustness of our sample grouping by bootstrapping, selecting 10 random genes 10,000 times, and counting the number of times this exact separation was achieved – which was 550 times.

### Intra-cluster WGCNA

To agnostically identify gene-gene relatedness within cluster 11, we performed WGCNA as described (see *Pseudo-bulk WGCNA*), selecting only the 5000 most variable genes using Seurat’s *FindVariableFeatures* and variance stabilising transformation. The scale free topology threshold utilised was 6, as per the ‘power estimate’. P-values were corrected for multiple testing as before.

### Module Preservation Analysis

To analyse the preservation of the relatedness of *C11orf53-*correlated 11q23.1 trans-eQTL targets, we defined an auxiliary module, comprised of the 12 trans-eQTL targets correlated with *C11orf53* expression (cor>0.5, p<0.05) within the ‘cluster 11 black’ module. Pseudo-bulked expression of each cluster was calculated and subset for the 5000 most variable genes as before, see *Pseudo-bulk WGCNA* and *Intra-cluster WGCNA*. The expression from each cluster was then raised to the same scale free topology threshold as cluster 11. The preservation of all modules, including the auxiliary module, was calculated in a pairwise manner with cluster 11, following the tutorial by Langfelder & Horvath (2008) (https://horvath.genetics.ucla.edu/html/CoexpressionNetwork/ModulePreservation/Tutorials/). To summarise module preservation results, we extracted cor.kME values – a measure of the similarity of auxiliary module genes with the auxiliary module eigengene, and equivalent results from cluster 11. We also extracted the nominal p-value for this correlation (log.p.cor.kME), which was subsequently un-logged. The mean connectivity of auxiliary module genes with the module eigengene was also calculated within each cluster. Pairwise correlations (p<0.05) between the expression of the auxiliary module genes with one another and *C11orf53* expression were plotted using corrplot (Wei & Simko, 2017).

### Linear modelling of cluster abundance and pseudo-bulked gene expression

Univariate linear modelling of cluster 11 abundance was performed on the TMM normalised, non-z-scored pseudo-bulked expression matrix, so that results are more relevant to further study. We fitted a linear model to all genes and performed empirical Bayes moderation using *limma* v3.46.0 (Ritchie et al., 2015). P-values were adjusted by Benjamini-Hochberg multiple testing correction (Benjamini & Hochberg, 1995).

### Differential abundance testing

Differential abundance testing was performed using miloR v0.99.8 (Dann et al., 2021). To mitigate any potential for package-specific artefacts of the analysis, we first re-generated the k nearest neighbour graph using 250 nearest neighbours from the integrated expression. The graph was then built using 4 PCA components. Differential abundance testing was carried out using a Quasi-Likelihood F-test on TMM normalised cell proportions. Differentially abundant neighbourhood results were then annotated for their majority cluster proportion and those comprised of fewer than 80% of the majority cluster were removed.

## Author contributions

Conceptualisation: BTH, VR, MGD, SMF

Methodology: BTH, VR, SMF, MGD, AO, JPB, KD, MT, PGVS

Investigation: BTH

Original draft writing: BTH

Review & Editing: BTH, VR, JPB, KD, MT, AO, MGD, SMF

Funding Acquisition: SMF, MGD, FVND

Supervision: VR, MGD, SMF

## Supplementary Material

**Supplementary Table 1.** Smillie *et al*., (2019) patient metadata

| Subject ID | Status | Location | Gender |
| --- | --- | --- | --- |
| N8 | Healthy | #N/A | Female |
| N10 | Healthy | Right_Colon | Female |
| N11 | Healthy | Right_Colon | Male |
| N13 | Healthy | Right_Colon | Female |
| N15 | Healthy | Right_Colon | Male |
| N16 | Healthy | Right_Colon | Male |
| N17 | Healthy | Transverse_Colon | Male |
| N18 | Healthy | Transverse_Colon | Female |
| N20 | Healthy | #N/A | Female |
| N21 | Healthy | Left_Colon | Female |
| N46 | Healthy | Right_Colon | Male |

**Supplementary Figure 1.**
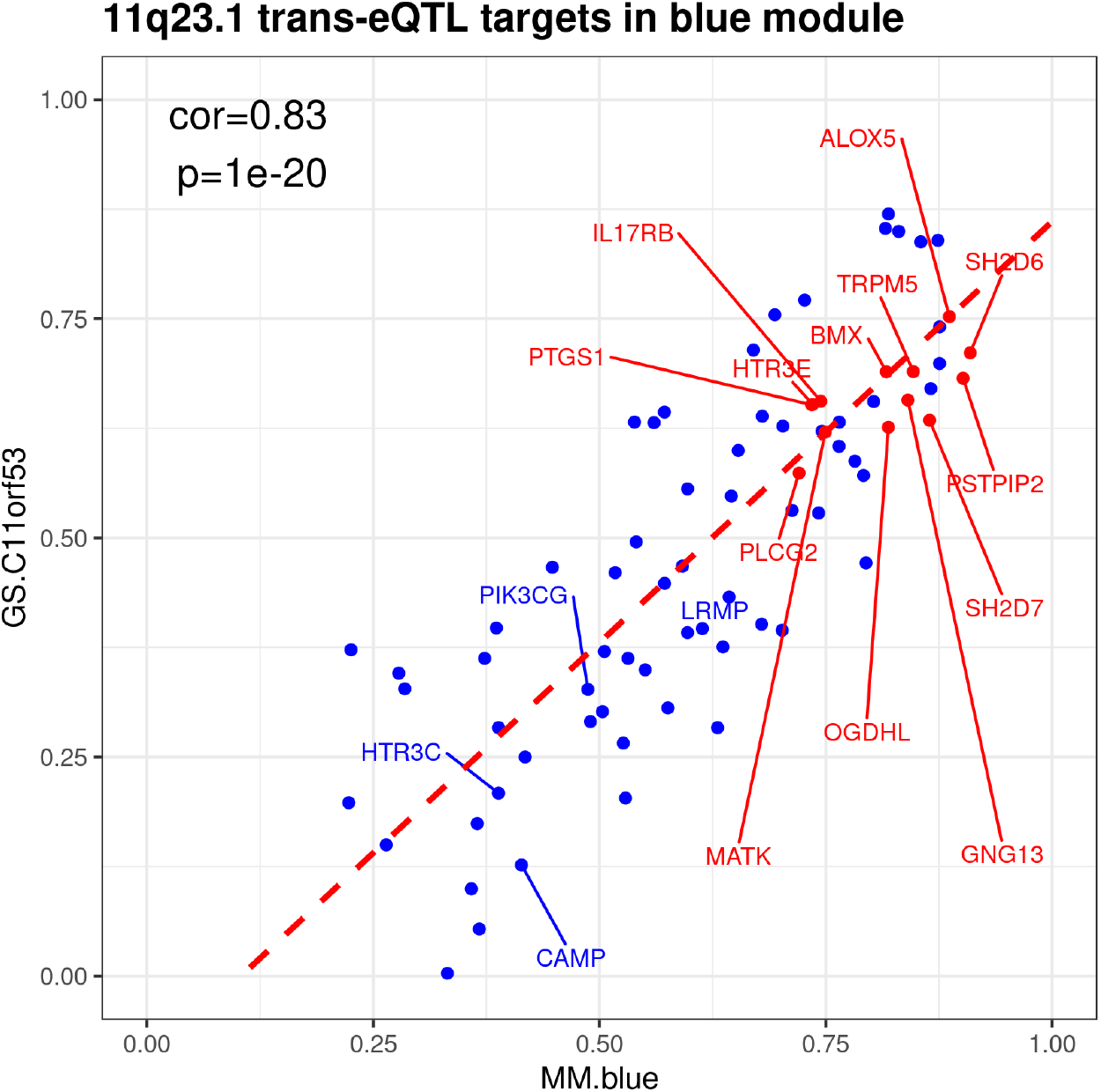
Correlation of Module Membership (MM) and Gene Significance (GS) for *C11orf53* expression in the blue module. 11q23.1 trans-eQTL targets are highlighted, red if MM.blue and GS.C11orf53 > 0.5.

**Supplementary Table 2.** Blue module genes, corresponding Gene Significance for *C11orf53* (GS.C11orf53) and p-value of Gene Significance for *C11orf53* (p.GS.C11orf53). Red=11q23.1 trans-eQTL target (Vaughan-Shaw *et al*., 2021)

| Gene_symbol | GS.C11orf53 | p.GS.C11orf53 |  |  |  |
| --- | --- | --- | --- | --- | --- |
| *MFSD7* | 0.869516061 | 1.49E-07 | *IKZF4* | 0.528361091 | 0.011477883 |
| *P2RY2* | 0.852996421 | 4.57E-07 | *FAM65A* | 0.495417788 | 0.019052761 |
| *FANK1* | 0.849691779 | 5.63E-07 | *GFI1B* | 0.471411163 | 0.026778703 |
| *LINC00261* | 0.839485426 | 1.04E-06 | *SGK223* | 0.467790056 | 0.028134237 |
| *KLK4* | 0.837697653 | 1.15E-06 | *RNF144A* | 0.466520671 | 0.028622119 |
| *PLEKHG4* | 0.771254176 | 2.65E-05 | *CHSY1* | 0.460356935 | 0.031087422 |
| *RNF32* | 0.754523148 | 4.97E-05 | *DKK3* | 0.448035289 | 0.036517806 |
| *ALOX5* | 0.752415106 | 5.36E-05 | *ZFP1* | 0.432558977 | 0.044363985 |
| *PTPRD* | 0.740741398 | 8.05E-05 | *AFAP1L2* | 0.401466056 | 0.064037752 |
| *ASTE1* | 0.714297945 | 0.000188259 | *TBC1D24* | 0.397206607 | 0.067180992 |
| *SH2D6* | 0.71110102 | 2.07E-04 | *LRMP* | 0.396518404 | 0.067699644 |
| *HCK* | 0.698902984 | 2.96E-04 | *PODXL* | 0.394606685 | 0.069156315 |
| *TRPM5* | 0.689672239 | 3.84E-04 | *SLC25A30* | 0.391914382 | 0.071247869 |
| *BMX* | 0.689502913 | 3.85E-04 | *ARHGEF3* | 0.375600915 | 0.084958526 |
| *PSTPIP2* | 0.682088126 | 4.71E-04 | *ARHGEF2* | 0.372293733 | 0.087962678 |
| *MYOM1* | 0.670290717 | 6.42E-04 | *FILIP1L* | 0.370412873 | 0.089706055 |
| *GNG13* | 0.657207652 | 8.90E-04 | *C20orf194* | 0.362507448 | 0.097315042 |
| *IL17RB* | 0.655879017 | 0.000918972 | *NOTCH4* | 0.362506253 | 0.097316226 |
| *CYR61* | 0.655513165 | 0.000927169 | *ARHGAP31* | 0.3494408 | 0.11091913 |
| *HTR3E* | 0.652281 | 0.001002333 | *PGAM4* | 0.34556863 | 0.115204208 |
| *PTGS1* | 0.651790925 | 0.001014171 | *FGFRL1* | 0.32779629 | 0.136419162 |
| *WWP2* | 0.643465569 | 0.001234253 | *PIK3CG* | 0.327080559 | 0.137328021 |
| *FYB* | 0.638720231 | 0.001376965 | *ATF3* | 0.305953301 | 0.16612758 |
| *SH2D7* | 0.63416094 | 0.001527043 | *PPP1R3B* | 0.301754652 | 0.172314946 |
| *SNRPA* | 0.632098175 | 0.00159937 | *CAPN9* | 0.290378683 | 0.189872811 |
| *AC025335.1* | 0.632010561 | 0.001602506 | *APH1B* | 0.283453554 | 0.201136526 |
| *RAI1* | 0.631424527 | 0.001623611 | *MT1H* | 0.283442955 | 0.201154103 |
| *MARCKSL1* | 0.627445473 | 0.001773235 | *GNB5* | 0.265793153 | 0.23186717 |
| *OGDHL* | 0.626140466 | 0.00182478 | *SNORA31* | 0.249996893 | 0.261829995 |
| *CRYM* | 0.621805521 | 0.002005201 | *HTR3C* | 0.20872 | 0.351252864 |
| *MATK* | 0.620641028 | 0.002056158 | *TRIB1* | 0.203124291 | 0.364604113 |
| *ALOX5AP* | 0.604524135 | 0.002881786 | *COL1A1* | 0.197820332 | 0.377525123 |
| *RP11-315O6.1* | 0.599692821 | 0.003177792 | *PIP5KL1* | 0.174089375 | 0.438438491 |
| *MSLN* | 0.587571695 | 0.004034646 | *CGNL1* | 0.149829973 | 0.505720737 |
| *PLCG2* | 0.573987877 | 0.005216381 | *CAMP* | 0.126754067 | 0.574047366 |
| *SCRN1* | 0.571174034 | 0.005494069 | *ISPD* | 0.099579334 | 0.659282892 |
| *CD96* | 0.555950868 | 0.007218247 | *CAPRIN2* | 0.053739047 | 0.812256115 |
| *L3MBTL1* | 0.547690051 | 0.008327128 | *DLG5-AS1* | 0.003234775 | 0.988601201 |
| *RP11-1220K2.2* | 0.531364238 | 0.010932727 |  |  |  |

**Supplementary Figure 2.**
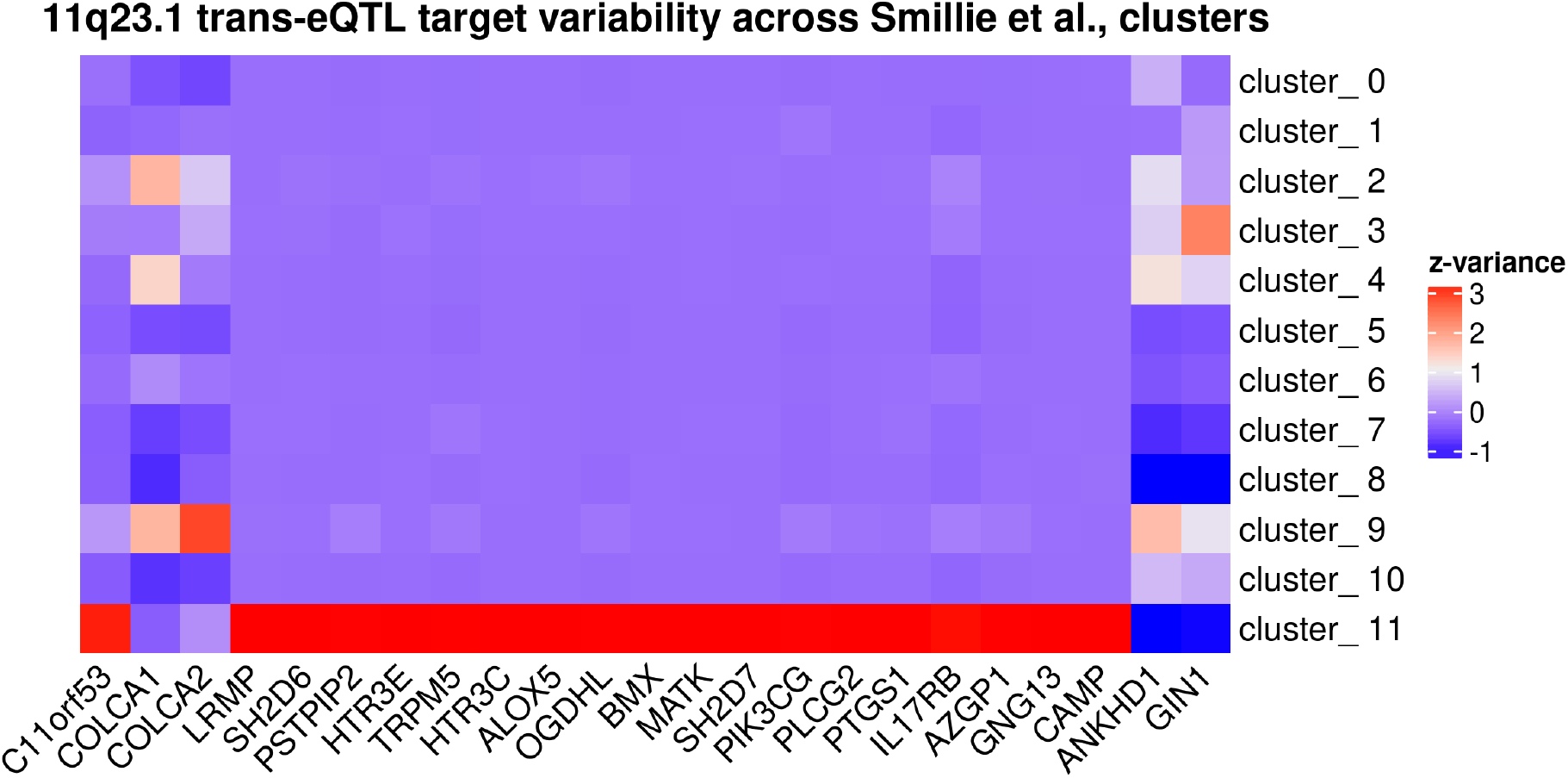
Investigating 11q23.1 eQTL target variance at the single cell level. Variance of 11q23.1 eQTL targets across clusters at the single cell level in the Smillie *et al*., (2019) data. Variance is z-scored within genes.

**Supplementary Figure 3.**
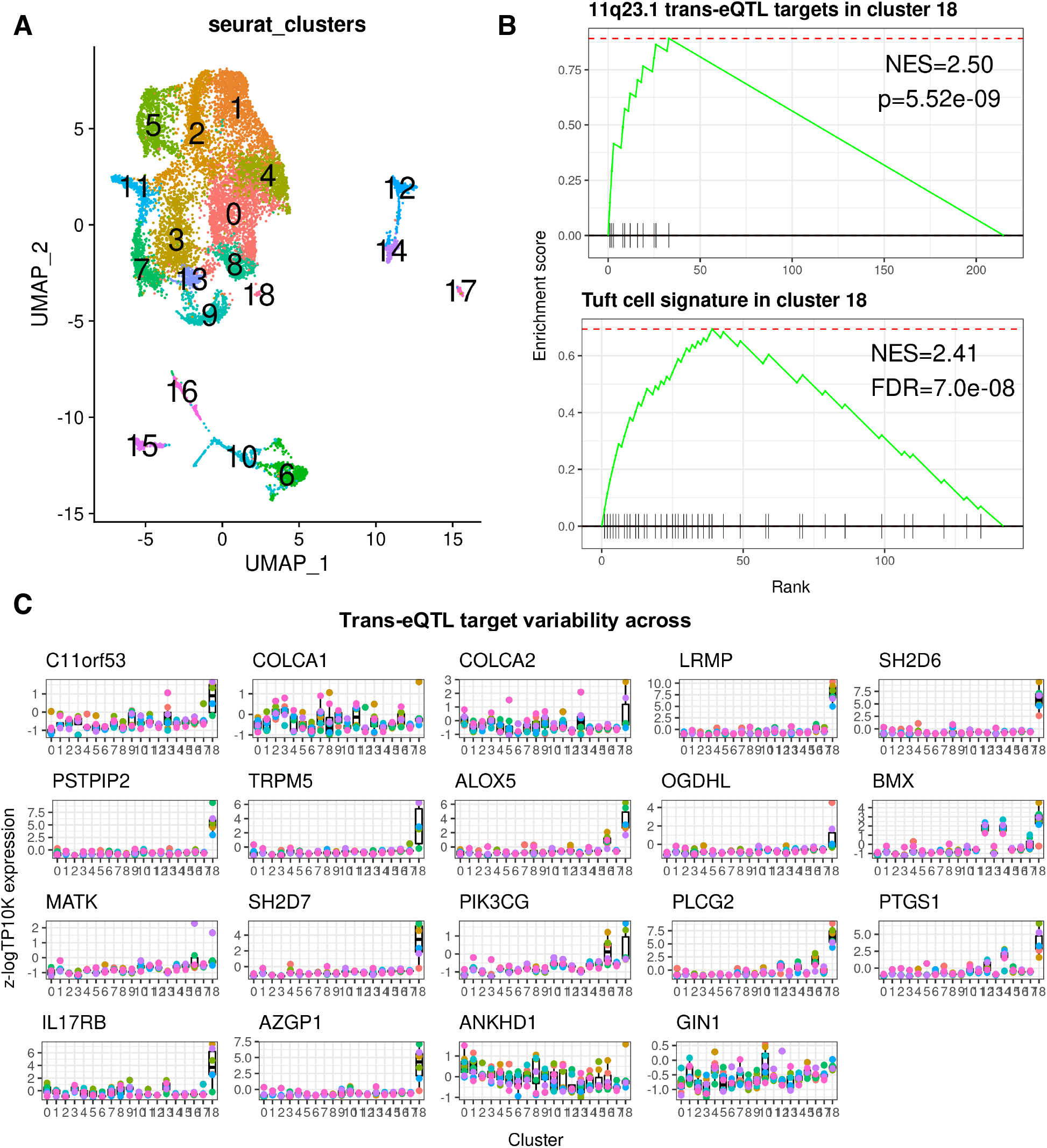
Replication of 11q23.1 eQTL expression mapping in independent dataset. (A) UMAP of 11,651 healthy colonic epithelial scRNASeq (Elmentaite *et al*., 2021). (B) Upper: GSEA of 11q23.1 trans-eQTLs (Vaughan-Shaw *et al*., 2021; FDR<0.05) in cluster 18. Lower: GSEA of putative colonic tuft cell signature (Smillie *et al*., 2019) in cluster 18. (D) Relative expression of 11q23.1 trans-eQTLs (FDR<0.05) across clusters.

**Supplementary Table 3.** 11q23.1 trans-eQTL targets identified as cluster 18 markers from Elmentaite *et al*., (2021). Markers calculated using MAST (McDavid *et al*., 2020). Avg_log2FC=average log2 fold change between cluster 11 and all other clusters. Pct1=Proportion of cluster 18 cells expressing gene. Pct2=Proportion of non-cluster 18 cells expressing gene.

| Gene | p_val | avg_log2FC | pct.1 | pct.2 | p_val_adj |
| --- | --- | --- | --- | --- | --- |
| *LRMP* | 2.04E-73 | 2.614295875 | 0.903 | 0.004 | 3.19E-69 |
| *IL17RB* | 6.17E-69 | 1.454372429 | 0.516 | 0.033 | 9.64E-65 |
| *SH2D6* | 3.81E-64 | 2.521466548 | 0.806 | 0.002 | 5.95E-60 |
| *PLCG2* | 1.11E-50 | 2.182226523 | 0.71 | 0.011 | 1.73E-46 |
| *PSTPIP2* | 1.31E-45 | 1.656282707 | 0.581 | 0.006 | 2.04E-41 |
| *TRPM5* | 6.51E-44 | 1.403600676 | 0.581 | 0.002 | 1.02E-39 |
| *SH2D7* | 2.90E-39 | 1.090698798 | 0.484 | 0.001 | 4.53E-35 |
| *AZGP1* | 9.92E-39 | 1.061625996 | 0.452 | 0.005 | 1.55E-34 |
| *PTGS1* | 8.37E-30 | 1.282060245 | 0.516 | 0.01 | 1.31E-25 |
| *ALOX5* | 4.14E-29 | 1.32717779 | 0.452 | 0.004 | 6.46E-25 |
| *BMX* | 1.08E-22 | 1.007016149 | 0.419 | 0.012 | 1.68E-18 |

**Supplementary Figure 4.**
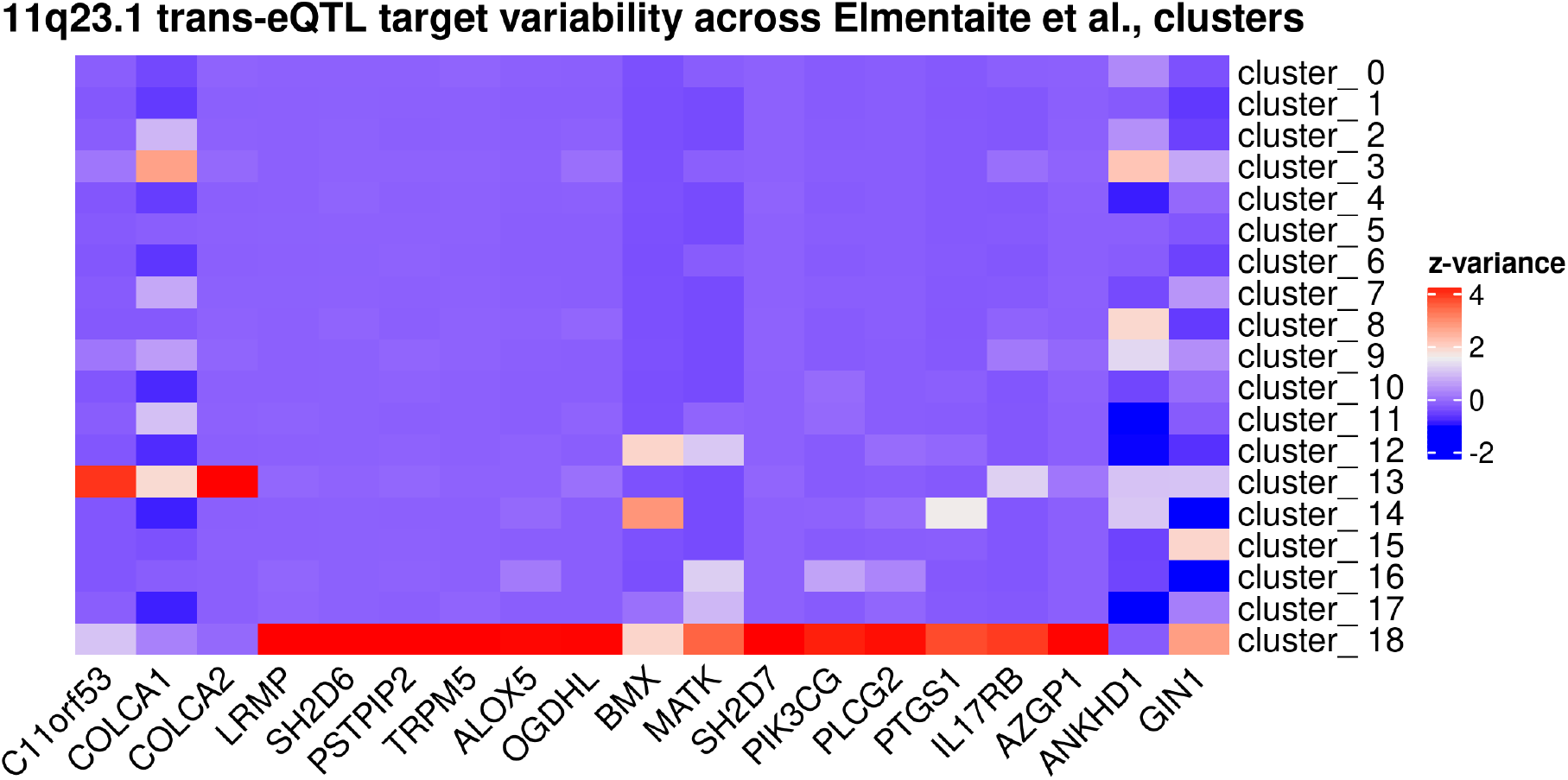
Investigating 11q23.1 eQTL target variance at the single cell level. Variance of 11q23.1 eQTL targets across clusters at the single cell level in Elmentaite *et al*., (2021) data. Variance is z-scored within genes.

**Supplementary Table 4.** Elmentaite *et al*., (2019) samples are homozygous non-risk at rs3087967. rs3087967 genotype results of both freebayes (Garrison et al., 2012) and Bcftools (Danecek et al., 2021) variant calling methods. T=colorectal cancer risk allele, C=non-risk allele

| Sample | Subject | freebayes_rs3087967 | samtools_rs3087967 |
| --- | --- | --- | --- |
| Human_colon_16S8123908 | 411C | CC | CC |
| Human_colon_16S8000473 | 417C | T/C | CC |
| Human_colon_16S8000477 | 417C | CC | CC |
| Human_colon_16S8000481 | 417C | CC | CC |
| Human_colon_16S8000479 | 417C | CC | CC |
| Human_colon_16S8000475 | 417C | CC | CC |
| Human_colon_16S8002627 | 432c | CC | CC |
| Human_colon_16S8002630 | 432c | CC | CC |

**Supplementary Table 5.** Variability of the majority of 11q23.1 trans-eQTL targets is greater in Smillie *et al*., (2019) cluster 11 compared with Elmentaite *et al*., (2021) cluster 18. Variability analysed within single cells using Seurat *FindVariableGenes* (Hao et al., 2021). FC = Fold change of variability in Smillie *et al*., cluster 11 compared with Elmentaite *et al*., (2021) cluster 18

| Gene | Smillie_cluster11 | Elmentaite_cluster18 | FC |
| --- | --- | --- | --- |
| *C11orf53* | 1.275873075 | 0.656341714 | 1.94391587 |
| *COLCA1* | 1.346177989 | NA | NA |
| *COLCA2* | 1.381191066 | 0.834903033 | 1.65431315 |
| *LRMP* | 1.695513426 | 0.173077239 | 9.79628192 |
| *SH2D6* | 0.518050076 | 0.196087378 | 2.64193484 |
| *PSTPIP2* | 0.961252255 | 0.332407278 | 2.89179064 |
| *HTR3C* | 1.850394057 | NA | NA |
| *ALOX5* | 0.855502479 | 0.467755885 | 1.82895075 |
| *OGDHL* | 0.83318323 | 1.141132077 | 0.73013742 |
| *MATK* | 0.815029157 | NA | NA |
| *PIK3CG* | 1.103714386 | 0.73167938 | 1.50846726 |
| *PLCG2* | 0.871942447 | 0.249476603 | 3.49508706 |
| *PTGS1* | 0.673534891 | 0.461933218 | 1.45807849 |
| *IL17RB* | 0.8969865 | 0.467309219 | 1.91947101 |
| *GNG13* | 1.024981369 | NA | NA |
| *CAMP* | 2.260972819 | NA | NA |
| *ANKHD1* | 0.897739169 | NA | NA |
| *GIN1* | 1 | NA | NA |
| *SH2D7* | 0.658046427 | 0.328744382 | 2.00169634 |
| *BMX* | 1.360003183 | 0.360535132 | 3.77217936 |
| *HTR3E* | 0.848772333 | NA | NA |
| *AZGP1* | 1.029538915 | 0.515296922 | 1.99795277 |
| *TRPM5* | 0.599018856 | 0.314108919 | 1.90704186 |

**Supplementary Figure 5.**
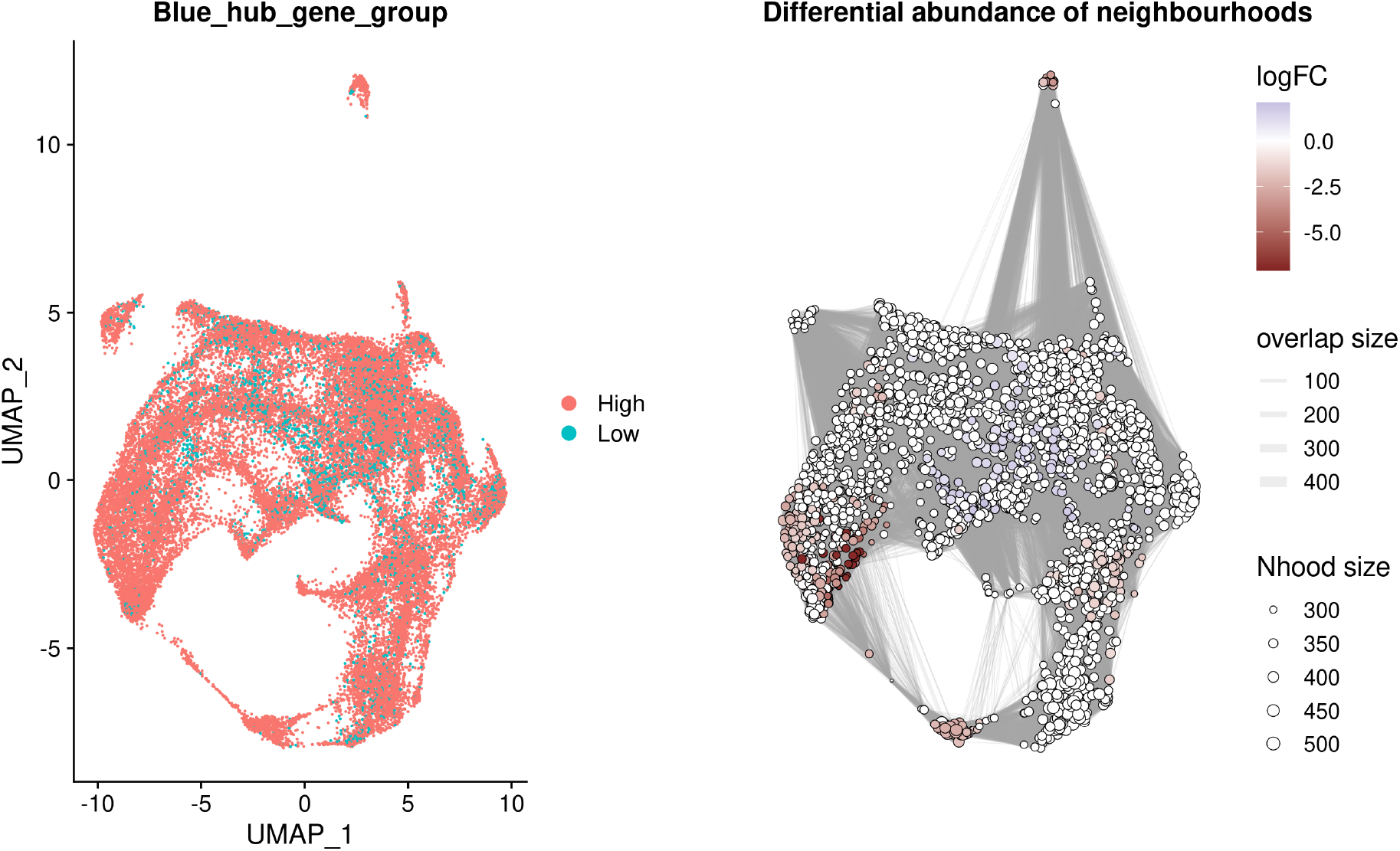
UMAP of cells from each Blue module hub gene group (left) and the differential abundance score of miloR-identified (Dann *et al*., 2021) neighbourhoods (right).

## Data availability

All data used in this study has been referenced within the Materials and Methods.

## Open Access

For the purpose of open access, the author has applied a Creative Commons Attribution (CC BY) licence to any Author Accepted Manuscript version arising from this submission.

